# Deriving robust biomarkers from multi-site resting-state data: An Autism-based example

**DOI:** 10.1101/075853

**Authors:** Alexandre Abraham, Michael Milham, Adriana Di Martino, R. Cameron Craddock, Dimitris Samaras, Bertrand Thirion, Gael Varoquaux

## Abstract

Resting-state functional Magnetic Resonance Imaging (R-fMRI) holds the promise to reveal functional biomarkers of neuropsychiatric disorders. However, extracting such biomarkers is challenging for complex multi-faceted neuropathologies, such as autism spectrum disorders. Large multi-site datasets increase sample sizes to compensate for this complexity, at the cost of uncontrolled heterogeneity. This heterogeneity raises new challenges, akin to those face in realistic diagnostic applications. Here, we demonstrate the feasibility of inter-site classification of neuropsychiatric status, with an application to the Autism Brain Imaging Data Exchange (ABIDE) database, a large (N=871) multi-site autism dataset. For this purpose, we investigate pipelines that extract the most predictive biomarkers from the data. These R-fMRI pipelines build participant-specific connectomes from functionally-defined brain areas. Connectomes are then compared across participants to learn patterns of connectivity that differentiate typical controls from individuals with autism. We predict this neuropsychiatric status for participants from the same acquisition sites or different, unseen, ones. Good choices of methods for the various steps of the pipeline lead to 67% prediction accuracy on the full ABIDE data, which is significantly better than previously reported results. We perform extensive validation on multiple subsets of the data defined by different inclusion criteria. These enables detailed analysis of the factors contributing to successful connectome-based prediction. First, prediction accuracy improves as we include more subjects, up to the maximum amount of subjects available. Second, the definition of functional brain areas is of paramount importance for biomarker discovery: brain areas extracted from large R-fMRI datasets outperform reference atlases in the classification tasks.

## 1. Introduction

In psychiatry, as in other fields of medicine, both the standardized observation of signs, as well as the symptom profile are critical for diagnosis. However, compared to other fields of medicine, psychiatry lacks accompanying objective markers that could lead to more refined diagnoses and targeted treatment [1]. Advances in non-invasive brain imaging techniques and analyses (*e. g.* [2, 3]) are showing great promise for uncovering patterns of brain structure and function that can be used as objective measures of mental illness. Such *neurophenotypes* are important for clinical applications such as disease staging, determination of risk prognosis, prediction and monitoring of treatment response, and aid towards diagnosis (*e. g.* [4]).

Among the many imaging techniques available, resting-state fMRI (R-fMRI) is a promising candidate to define functional neurophenotypes [5, 3]. In particular, it is non-invasive and, unlike conventional task-based fMRI, it does not require a constrained experimental setup nor the active and focused participation of the subject. It has been proven to capture interactions between brain regions that may lead to neuropathology diagnostic biomarkers [6]. Numerous studies have linked variations in brain functional architecture measured from R-fMRI to behavioral traits and mental health conditions such as Alzheimer disease (*e. g.* [7, 8], Schizophrenia (*e. g.* [9, 10, 11, 12]), ADHD, autism (*e. g.* [13]) and others (*e. g.* [14]). Extending these findings, predictive modeling approaches have revealed patterns of brain functional connectivity that could serve as biomarkers for classifying depression (*e. g.* [15]), ADHD (*e. g.* [16]), autism (*e. g.* [17]), and even age [18]. This growing number of studies has shown the feasibility of using R-fMRI to identify biomarkers. However questions about the readiness of R-fMRI to detect clinically useful biomarkers remain [13]. In particular, the reproducibility and generalizability of these approaches in research or clinical settings are debatable. Given the modest sample size of most R-fMRI studies, the effect of cross-study differences in data acquisition, image processing, and sampling strategies [19, 20, 21] has not been quantified.

Using larger datasets is commonly cited as a solution to challenges in reproducibility and statistical power [22]. They are considered a prerequisite to R-fMRI-based classifiers for the detection of psychiatric illness. Recent efforts have accelerated the generation of large databases through sharing and aggregating independent data samples[23, 24, 25]. However, a number of concerns must be addressed before accepting the utility of this approach. Most notably, the many potential sources of uncontrolled variation that can exist across studies and sites, which range from MRI acquisition protocols (*e. g.* scanner type, imaging sequence, see [26]), to participant instructions (*e. g.* eyes open vs. closed, see [27]), to recruitment strategies (age-group, IQ-range, level of impairment, treatment history and acceptable comorbidities). Such variation in aggregate samples is often viewed as dissuasive, as its effect on diagnosis and biomarker extraction is unknown. It commonly motivates researchers to limit the number of sites included in their analyses at the cost of sample size.

Cross-validated results obtained from predictive models are more robust to inhomogeneities: they measure model generalizability by applying it to unseen data, *i. e.*, data not used to train the model. In particular, leave-out cross-validation strategies, which remove single individuals (or random subsets), are common in biomarkers studies. However, these strategies do not measure the effect of potential site-specific confounds. In the present study we leverage aggregated R-fMRI samples to address this problem. Instead of leaving out random subsamples as test sets, we left out entire sites to measure performance in the presence of uncontrolled variability.

Beyond challenges due to inter-site data heterogeneity, choices in the functional-connectivity data-processing pipeline further add to the variability of results [28, 27, 29]. While preprocessing procedures are now standard, the different steps of the prediction pipeline vary from one study to another. These entail specifying regions of interest, extracting regional time courses, computing connectivity between regions, and identifying connections that relate to subject’s phenotypes [15, 30, 31, 32].

Lack of ground truth for brain functional architecture undermines the validation of R-fMRI data-processing pipelines. The use of functional connectivity for individual prediction suggests a natural figure of merit: prediction accuracy. We contribute quantitative evaluations, to help settling down on a parameter-free pipeline for R-fMRI. Using efficient implementations, we were able to evaluate many pipeline options and select the best method to estimate atlases, extract connectivity matrices, and predict phenotypes.

To demonstrate that pipelines to extract R-fMRI neuro-phenotypes can reliably learn inter-site biomarkers of psychiatric status on inhomogeneous data, we analyzed R-fMRI in the Autism Brain Imaging Data Exchange (ABIDE) [25]. It compiles a dataset of 1112 R-fMRI participants by gathering data from 17 different sites. After preprocessing, we selected 871 to meet quality criteria for MRI and phenotypic information. Our inter-site prediction methodology reproduced conditions found under most clinical settings, by leaving out whole sites and using them as newly seen test sets. To validate the robustness of our approach, we performed nested cross-validation and varied samples per inclusion criteria (*e. g.* sex, age). Finally, to assess losses in predictive power associated with using a heterogeneous aggregate dataset instead of uniformly defined samples, we included a comparison of intra- and inter-site prediction strategies.

## 2. Material and methods

A connectome is a functional-connectivity matrix between a set brain regions of interest (ROIs). We call such a set of ROIs an atlas, even though some of the methods we consider extract the regions from the data rather than relying on a reference atlas (see Figure 1). We investigate here pipelines that discriminate individuals based on the connection strength of this connectome [33], with a classifier on the edge weights [15].

**Figure 1:**
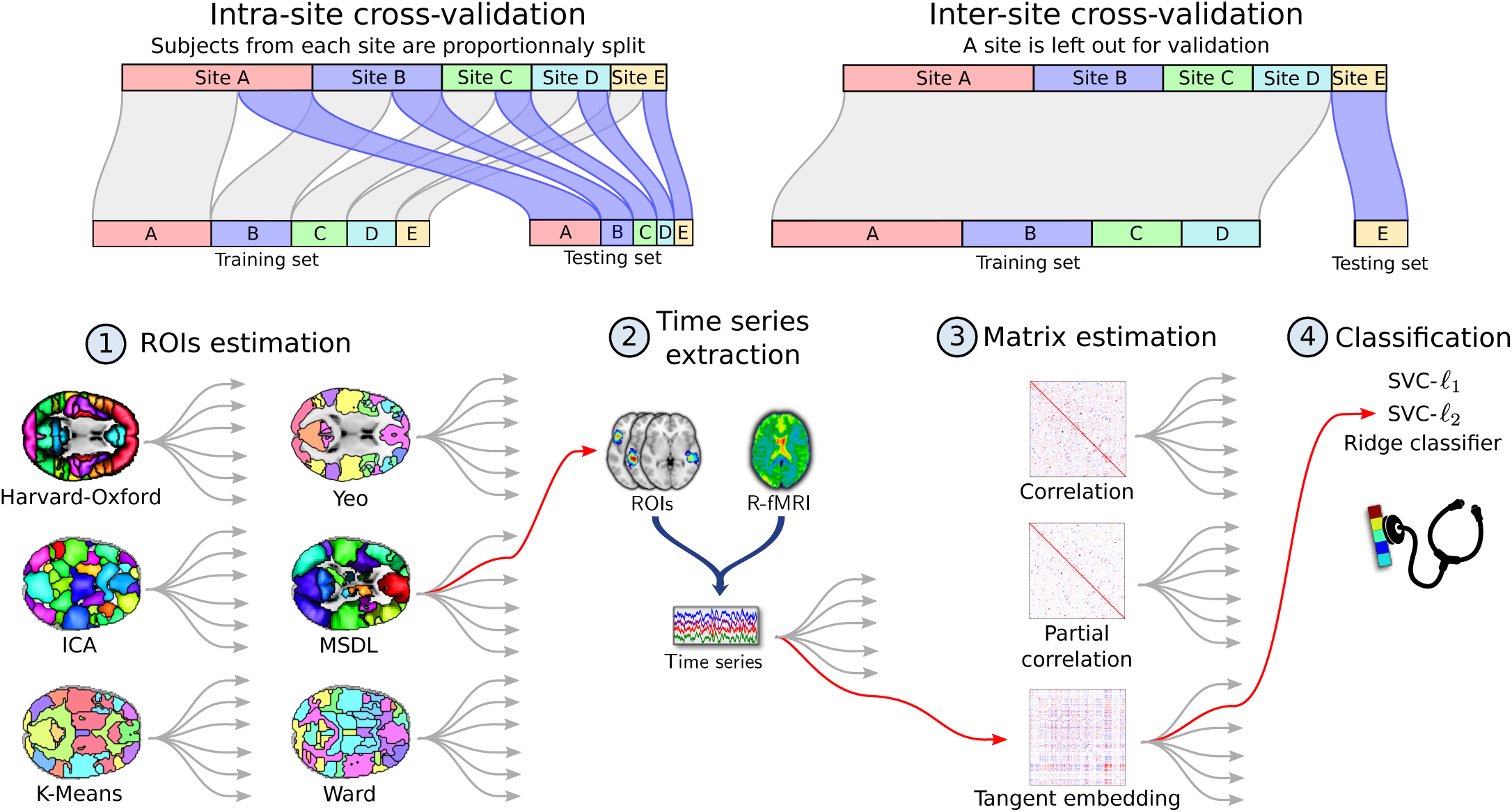
Functional MRI analysis pipeline. Cross-validation schemes used to validate the pipeline are presented above. **Intra-site** cross-validation consists of randomly splitting the participants into training and testing sets while preserving the ratio of samples for each site and condition. **Inter-site** cross-validation consists of leaving out participants from an entire site as testing set. In the first step of the pipeline, regions of interest are estimated from the training set. The second step consists of extracting signals of interest from all the participants, which are turned into connectivity features via covariance estimation at the third step. These features are used in the fourth step to perform a supervised learning task and yield an accuracy score. An example of pipeline is highlighted in red. This pipeline is the one that gives best results for inter-site prediction. Each model is decribed in the section relative to material and methods.

Specifically, we consider connectome-extraction pipelines composed of four steps: 1) region estimation,2) time series extraction, 3) matrix estimation, and 4) classification –see Figure 1. We investigate different options for the various steps: brain parcellation method, connectivity matrix estimation, and final classifier.

In order to measure the impact of uncontrolled variability on prediction performance, we designed a diagnosis task where the model classifies participants coming from an acquisition site not seen during training.

### 2.1. Datasets: Preprocessing and Quality Assurance (QA)

We ran our analyses on the ABIDE repository [25], a sample aggregated across 17 independent sites. Since there was no prior coordination between sites, the scan and diagnostic/assessment protocols vary across sites ^1^. To easily replicate and extend our work, we rely on a publicly available preprocessed version of this dataset provided by the Preprocessed Connectome Project ^2^ initiative. We specifically used the data processed with the Configurable Pipeline for the Analysis of Connectomes [34] (C-PAC). Data were selected based on the results of quality visual inspection by three human experts who inspected for largely incomplete brain coverage, high movement peaks, ghosting and other scanner artifacts. This yielded 871 subjects out of the initial 1112.

Preprocessing included slice-timing correction, image realignment to correct for motion, and intensity normalization. Nuisance regression was performed to remove signal fluctuations induced by head motion, respiration, cardiac pulsation, and scanner drift [35, 36]. Head motion was modeled using 24 regressors derived from the parameters estimated during motion realignment [37], scanner drift was modeled using a quadratic and linear term, and physiological noise was modeled using the 5 principal components with highest variance from a decomposition of white matter and CSF voxel time series (CompCor) [38]. After regression of nuisance signals, fMRI was coregistered on the anatomical image with FSL BBreg, and the results where normalized to MNI space with the non-linear registration from ANTS [39]. Following time series extraction, data were detrended and standardized (dividing by the standard deviation in each voxel).

### 2.2. Step 1: Region definition

To reduce the dimensionality of the estimated connectomes, and to improve the interpretability of results, connectome nodes were defined from regions of interest (ROIs) rather than single voxels. ROIs can be either selected from a priori atlases or directly estimated from the correlation structure of the data being analyzed. Several choices for both approaches were used and compared to evaluate the impact of ROI definition on prediction accuracy. Note that the ROI were all defined on the training set to avoid possible overfitting.

We considered the following predefined atlases: **Harvard Oxford** [40], a structural atlas computed from 36 individuals’ T1 images, **Yeo** [41], a functional atlas composed of 17 networks derived from clustering [42] of functional connectivity on a thousand individuals, and **Craddock** [43], a multiscale functional atlas computed using constrained spectral clustering, to study the impact of the number of regions.

We derived data-driven atlases based on four strategies. We explored two clustering methods: **K-Means**, a technique to cluster fMRI time series [44, 45], which is a top-down approach minimizing cluster-level variance; and **Ward’s clustering**, which also minimizes a variance criterion using hierarchical agglomeration. Ward’s algorithm admits spatial constraints at no cost and has been extensively used to learn brain parcellations [45]. We also explored two decomposition methods: **Independent component analysis (ICA)** and **multi-subject dictionary learning (MSDL)**. ICA is a widely-used method for extracting brain maps from R-fMRI [46, 47] that maximizes the statistical independence between spatial maps. We specifically used the ICA implementation of [48]. MSDL extracts a group atlas based on dictionary learning and multi-subject modeling, and employs spatial penalization to promote contiguous regions [49]. Unlike MSDL and Ward clustering, ICA and K-Means do not take into account the spatial structure of the features and are not able to enforce a particular spatial constraint on their components. We indirectly provide such structure by applying a Gaussian smoothing on the data [45] with a full-width half maximum varying from 4mm to 12mm. Differences in the number of ROIs in the various atlases present a potential confound for our comparisons. As such, connectomes were constructed using only the 84 largest ROIs from each atlas, following [49]^3^.

### 2.3. Step 2: Time-series extraction

The common procedure to extract one representative time series per ROI is to take the average of the voxel signals in each region (for non overlapping ROIs) or the weighted average of non-overlapping regions (for fuzzy maps). For example, ICA and MSDL produce overlapping fuzzy maps. Each voxel’s intensity reflects the level of confidence that this voxel belongs to a specific region in the atlas. In this case, we found that extracting time courses using a spatial regression that models each volume of RfMRI data as a mixture of spatial patterns gives better results (details in Appendix A and Figure A3). For non-overlapping ROIs, this procedure is equivalent to weighted averaging on the ROI maps.

After this extraction, we remove at the region level the same nuisance regressors as at the voxel level in the preprocessing. Signals summarizing high-variance voxels (CompCor [38]) are regressed out: the 5 first principal components of the 2% voxels with highest variance. In addition, we extract the signals of ROIs corresponding to physiological or acquisition noise^4^ and regress them out to reject non-neural information [33]. Finally, we also regress out primary and secondary head motion artifacts using 24-regressors derived from the parameters estimated during motion realignment [37].

### 2.4. Step 3: Connectivity matrix estimation

Since scans in the ABIDE dataset do not include enough R-fMRI time points to reliably estimate the true covariance matrix for a given individual, we used the Ledoit-Wolf shrinkage estimator, a parameter-free regularized covariance estimator [50], to estimate relationships between ROI time series (details are given in Appendix B). For pipeline step 3, we studied three different connectivity measures to capture interactions between brain regions: *i)* the correlation itself, *ii)* partial correlations from the inverse covariance matrix [33, 51], and *iii)* the tangent embedding parameterization of the covariance matrix [48, 52]. We obtained one connectivity weight per pair of regions for each subject. At the group level, we ran a nuisance regression across the connectivity coefficients to remove the effects of site, age and gender.

### 2.5. Step 4: Supervised learning

As ABIDE represents a post-hoc aggregation of data from several different sites, the assessments used to measure autism severity vary somewhat across sites. As a consequence, autism severity scores are not directly quantitatively comparable between sites. However, the diagnosis between ASD and Typical Control (TC) is more reliable and has already been used in the context of classification [53, 13, 54].

Discriminating between ASD and TC individuals is a supervised-learning task. We use the connectivity measures between all pairs of regions, extracted in the previous step, as features to train a classifier across individuals to discriminate ASD patients from TC.

We explore different machine-learning methods^5^. We first rely on the most commonly used approach, the *ℓ*_2_-penalized support vector classification **SVC**. Given that we expect only a few connections to be important for diagnosis, we also include the *ℓ*_1_-penalized sparse **SVC**. Finally, we include **ridge regression**, which also uses an *ℓ*_2_ penality but is faster than the **SVC**. For all models we use the implementation of the scikit-learn library [55]^6^. We chose to work only with linear models for their interpretability. In particular, in subsection 3.7 we inspect classifier weights to identify which connections are the most important for prediction.

### 2.6. Exploring dataset heterogeneity

Previous studies of prediction using the ABIDE sample [13, 54] have included less than 25% of the available data, most likely to limit heterogeneity – not only due to differences in imaging, but also to factors such as age, sex, and handedness. We explored the effect of such choice on the results, by examining data subsets of different heterogeneity (see Table 1). These included: subsample #1 - **all participants** (871 individuals, 727 males and 144 females, 6 to 64 years old) is the full set of individuals that passed QA, subsample #2 - **largest sites** (736 individuals, 613 males and 123 females) 6 sites with less than 30 participants are removed from subsample #1, subsample #3 - **right handed males** (639 individuals) consists of the right-handed males from subsample #1, subsample #4 - **right handed males, 9–18 years** (420 individuals) is the restriction of subsample #3 to participants between 9 and 18 years old; subsample #5 - **right handed males, 9–18 yo, 3 sites** (226 individuals) are the individuals from subsample #4 belonging to the three largest sites (see Table A2 for more details)^7^.

**Table 1:** Subsets of ABIDE used in evaluation. We explore several subsets of ABIDE with different homogeneity.*Card*. stands for the cardinality of the dataset, *i. e.* the number of subjects. More details about these subsets are presented in the appendices -Table A2.

| Subsample | Card. | Sites | Selection criteria |
| --- | --- | --- | --- |
| 1 All subjects | 871 | 16 | All subjects after QC |
| 2 Biggest sites | 736 | 11 | Remove 6 smallest sites |
| 3 Right handed males | 639 | 14 | All subjects without left handed and women |
| 4 Right handed males, 9-18 yo | 420 | 14 | Right handed males between 9 and 18 years old |
| 5 Right handed males, 9-18 yo, 3 sites | 226 | 3 | Young right handed males from 3 biggest sites |

### 2.7. Experiments: prediction on heterogeneous data

ABIDE datasets come from 17 international sites with no prior coordination, which is a typical clinical application setting. In this setting, we want to: *i)* measure the performance of different prediction pipelines, *ii)* use these measures to find the best options for each pipeline step (Figure 1) and, *iii)* extract ASD biomarkers from the best pipeline.

*Cross-validation.* In order to measure the prediction accuracy of a pipeline, we use 10-fold cross-validation by training it exclusively on training data (including atlas estimation) and predicting on the testing set. Any hyperparameter of the various methods used inside a pipeline is set internally in a nested cross-validation.

We used two validation approaches. **Inter-site prediction** addresses the challenges associated with aggregate datasets by using whole sites as testing sets. This scenario also closely simulates the real world clinical challenge of not being able to sample data from every possible site. Note that this cross-validation can only be applied on a dataset with at least 10 acquisition sites, in order to leave out a different site in each fold. We do not apply it on subsample #5 that has been restricted to 3 sites to reduce sitespecific variability. **Intra-site prediction** builds training and testing sets are as homogeneous as possible, reducing site-related variability. It is based on stratified shuffle split cross-validation, which splits participants into training and test sets while preserving the ratio of samples for each site and condition. We used 80% of the dataset for training and the remaining for testing.

*Learning curves.* A learning curve measures the impact of the number of training samples on the performance of a predictive model. For a given prediction task, the learning curve is constructed by varying the number of samples in the training set with a constant test set. Typically, an increasing curve indicates that the model can still gain additional useful information from more data. It is expected that the performance of a model will plateau at some point.

*Summarizing prediction scores for a pipeline choice.* To quantify the impact of the different options on the prediction accuracy, we select representative prediction scores for each model. Indeed, for ICA, K-Means, and MSDL, we explore several values for each parameters and thus obtain a range of prediction values. For these methods, we use the 10% best-performing pipelines for post-hoc analysis ^8^. We do not rely only on the top-performing pipelines, as they may result from overfitting, nor the complete set of pipelines as some pipelines yield bad scores because of bad parametrization and may artificially lower the performance of these methods.

*Impact of pipeline choices on prediction accuracy.* In order to determine the best pipeline option at each step, we want to measure the impact of these choices on the prediction scores. In a full-factorial ANOVA, we then fit a linear model to explain prediction accuracy from the choice of pipeline steps, modeled as categorical variables. We report the corresponding effect size and its 95% confidence interval.

*Pairwise comparison of pipeline choice.* Two options can also be compared by comparing the scores between pairs of pipelines that differ only by that option using a Wilcoxon test that does not rely on Gaussian assumptions.

*Extracting biomarkers.* For a given pipeline, the final classifier yields a biomarker to diagnose between HC and ASD. To characterize this biomarker, we measure *p*-values on the classifier weights. We obtain the null distribution of the weights by permuting the prediction labels.

### 2.8. Computational aspects

For data-driven brain atlas models (*e. g.* ICA), region-extraction methods entail feature learning on up to 300 GB of data, too big to fit in memory. Systematic study requires many cross-validations and subsamples and thus careful computational optimizations. Hence, for K-Means and ICA, we reduced the length of the time series by applying PCA dimensionality reduction with efficient randomized SVD [57]. MSDL uses algorithmic optimizations [49] to process a large number of subjects in a reasonable time ^9^ (2 hours for 871 subjects). However, it is still the most costly method because of the number of runs required to find the optimal value for its 3 parameters.

For reproducibility, we rely on the publicly available implementations of the scikit-learn package [55] for all the clustering algorithms, connectivity matrix estimators and predictors.

## 3. Results

Here we present the most notable trends emerging from our analyses. More details are provided in supplementary materials. First, we compared inter- and intra-site prediction of ASD diagnostic labels while varying the number of subjects in the training set. Second, in a post-hoc analysis, we identified the pipeline choices most relevant for prediction and proposed a good choice of pipeline steps for prediction. Finally, by analyzing the weights of the classifiers, we highlighted functional connections that best discriminate between ASD and TC.

**Table 2:**
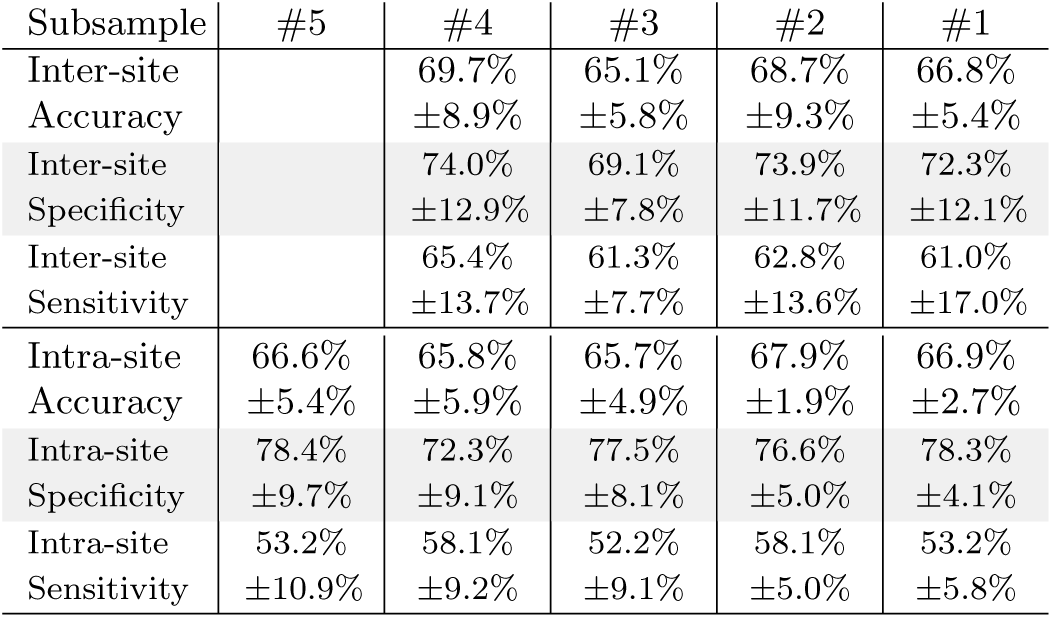
Average accuracy, specificity and sensitivity scores (and standard deviation) for top performing pipelines. Accuracy is the fraction of well-classified individuals (chance level is at 53.7%). Specificity is the fraction of well classified healthy individuals among all healthy individuals (percentage of well-classified negatives). Sensitivity is the ratio of ASD individuals among all subjects classified as ASD (percentage of positives that are true positives). Results per atlas are available in the appendices –Table A5.

| Subsample | #5 | #4 | #3 | #2 | #1 |
| --- | --- | --- | --- | --- | --- |
| Inter-site Accuracy |  | 69.7%<br>±8.9% | 65.1%<br>±5.8% | 68.7%<br>±9.3% | 66.8%<br>±5.4% |
| Inter-site Specificity |  | 74.0%<br>±12.9% | 69.1%<br>±7.8% | 73.9%<br>±11.7% | 72.3%<br>±12.1% |
| Inter-site Sensitivity |  | 65.4%<br>±13.7% | 61.3%<br>±7.7% | 62.8%<br>±13.6% | 61.0%<br>±17.0% |
| Intra-site Accuracy | 66.6%<br>±5.4% | 65.8%<br>±5.9% | 65.7%<br>±4.9% | 67.9%<br>±1.9% | 66.9%<br>±2.7% |
| Intra-site Specificity | 78.4%<br>±9.7% | 72.3%<br>±9.1% | 77.5%<br>±8.1% | 76.6%<br>±5.0% | 78.3%<br>±4.1% |
| Intra-site Sensitivity | 53.2%<br>±10.9% | 58.1%<br>±9.2% | 52.2%<br>±9.1% | 58.1%<br>±5.0% | 53.2%<br>±5.8% |

### 3.1. Overall prediction results

For inter-site prediction, the highest accuracy obtained on the whole dataset is 66.8% (see Table 2) which exceeds previously published ABIDE findings [53, 13, 54] and chance at 53.7%^10^. Importantly, performance increases steadily with training set size, for all subsamples (Figures 2 and A6). This increase implies that optimal classification performance is not reached even for the largest dataset tested.

**Figure 2:**
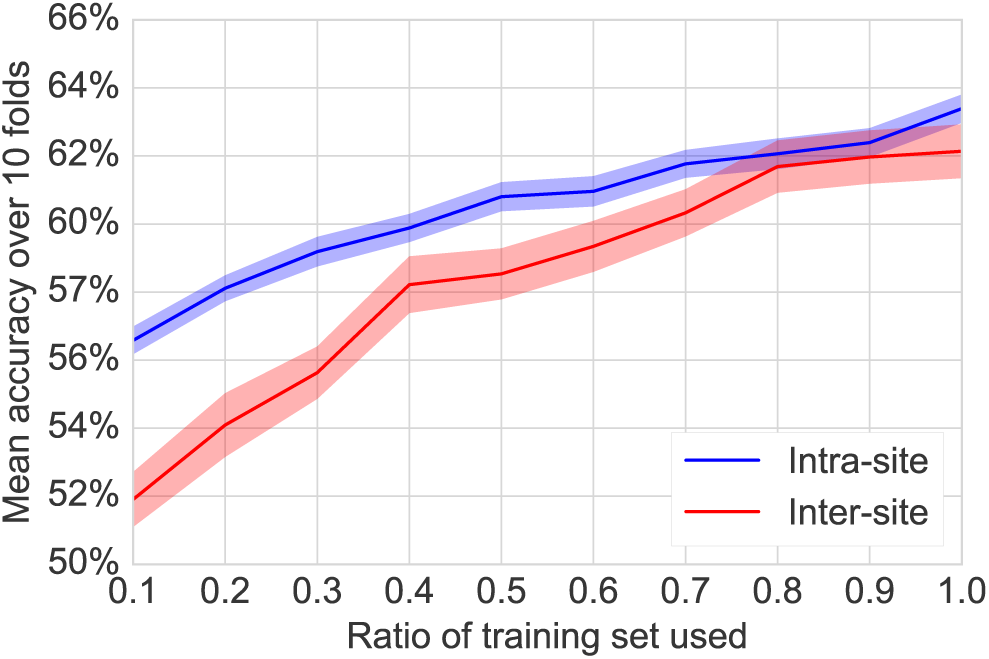
Learning curve. Classification results obtained by varying the ratio of the training set using to train the classifier while keeping a fixed testing set. The colored bands represent the standard error of the prediction. A score increasing with the number of subjects, *i. e.* a positive slope indicates that the addition of new subjects improves performance. This curve is an average of the results obtained on several subsamples. Detailed results per subsamples are presented in the appendices –Figure A6

The main difference between intra-site and inter-site settings is that the variability of inter-site prediction is higher (Table 2). However, this difference disappears with a large enough training set (Figure 2). This is encouraging for clinical applications.

As shown in Figure 3, the best predicting pipeline is MSDL, tangent space embedding and *l*_2_-regularized classifiers (SVC-*l*_2_ and ridge classifier). All effects are observed with *p* < 0.01. Detailed pairwise comparisons (Figure 4) confirm these trends by showing the effect of each option compared to the best ones. Results of intra-site prediction have smaller standard error, confirming higher variability of inter-site prediction results (Figure A5).

**Figure 3:**
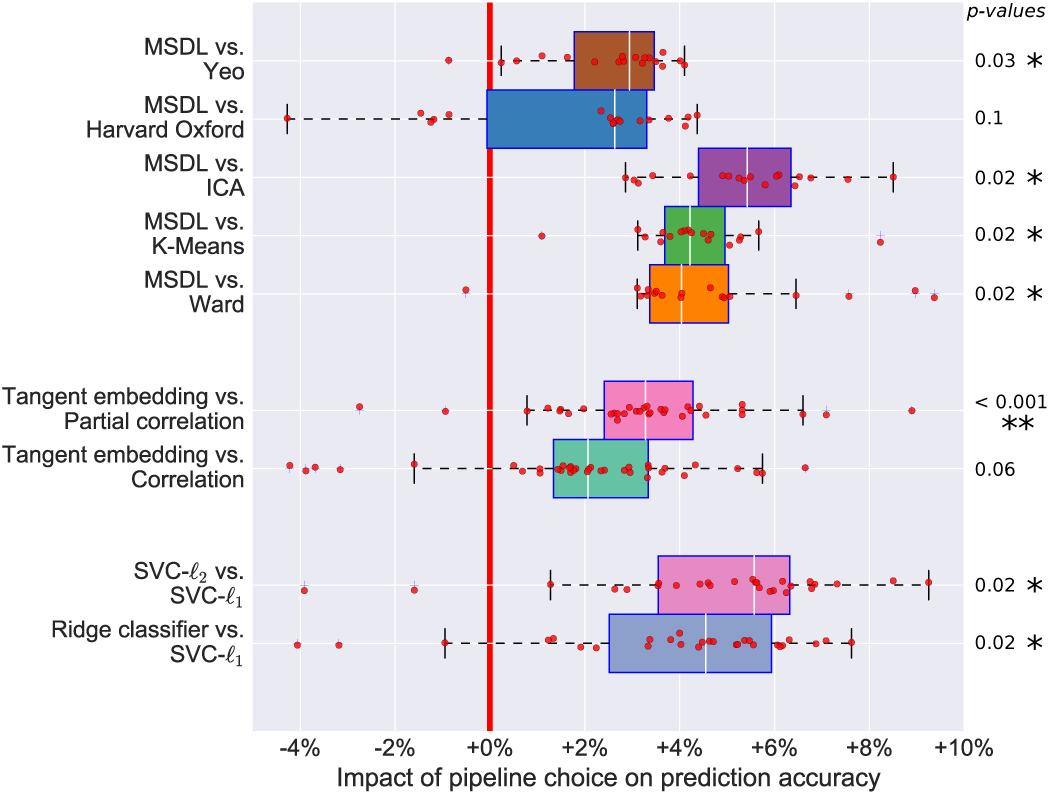
Impact of pipeline steps on prediction. Each block of bars represents a step of the pipeline (namely step 1, 3 and 4). We also prepended two steps corresponding to the cross-validation schemes and ABIDE subsamples. Each bar represents the impact of the corresponding option on the prediction accuracy, relative to the mean prediction. This effect is measured via a full-factorial analysis of variance (ANOVA), analyzing the contribution of each step in a linear model. Each step of the pipeline is considered as a categorical variable. Error bars give the 95% confidence interval. Multi Subject Dictionary Learning (MSDL) atlas extraction method gives significantly better results while reference atlases are slightly better than the mean. Among all matrix types, tangent embedding is the best on all ABIDE subsets. Finally, *l*_2_-regularized classifiers perform better than *l*_1_-regularized ones. Results for each subsamples are presented in the appendices –Figure A8.

**Figure 4:**
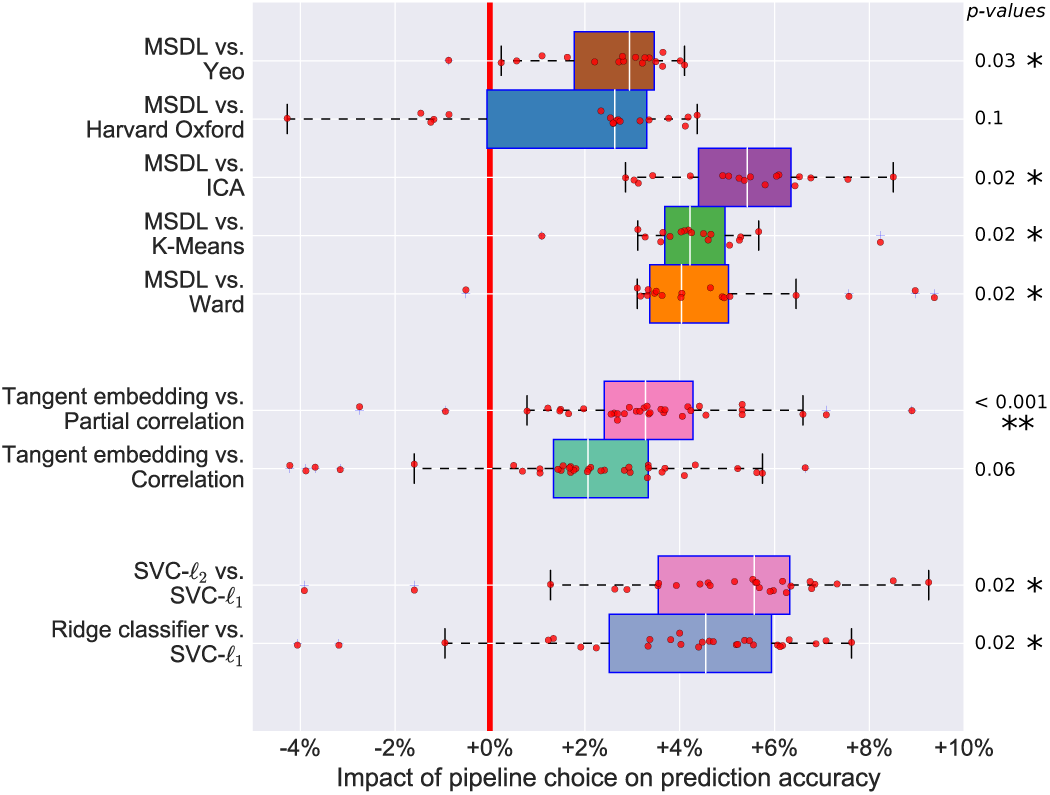
Comparison of pipeline options. Each plot compares the classification accuracy scores if one parameter is changed and the others kept fixed. We measured statistical signficance using a Wilcoxon signed rank test and corrected for multiple comparisons. Detailed one-to-one comparison of the scores are shown in the appendices –Figure A5

### 3.2. Effect of the choice of atlas

The choice of atlas appears to have the greatest impact on prediction accuracy (Figure 3). Estimation of an atlas from the data with MSDL, or using reference atlases leads to maximal performance. Of note, for smaller samples, data-driven atlas-extraction methods other than MSDL perform poorly (Figure A8). In this regime, noise can be problematic and limit the generalizability of atlases derived from data.

MSDL performs well regardless of sample size, and the performance gain is significant compared to all other strategies apart from the Harvard-Oxford atlas (Figure 4). This is likely attributable to MSDL’s strong spatially-structured regularization that increases its robustness to noise.

### 3.3. Effect of the covariance matrix estimator

While correlation and partial correlation approaches are currently the two dominant approaches to connectome estimation in the current R-fMRI literature, our results highlight the value of tangent-space projection that outperforms the other approaches in all settings (Figure 3 and Figure 4), though the difference is significant only compared with correlation matrices (Figure 4). This gain in prediction accuracy is not without a cost, as the matrices derived cannot be read as correlation matrices. Yet, the differences in connectivity that they capture can be directly interpreted [58].

Among the correlation-based approaches, full correlations outperformed partial correlations, most notably for intra-site prediction. These limitations of partial correlations may reflect estimation challenges^11^ with the relatively short time series included in ABIDE datasets (typically 5–6 minutes per participant).

### 3.4. Effect of the final classifier

The results show that *ℓ*_2_-regularized classifiers perform best in all settings (Figure 3 and Figure 4). This may be due to the fact that a global hypoconnectivity, as previously observed in ASD patients, is not well captured by *ℓ*_1_-regularized classifiers, which induce sparsity. In addition *ℓ*_2_-penalization is rotationally-invariant, which means that it is not sensitive to an arbitrary mixing of features. Such mixing may happen if a functional brain module sits astride several regions. Thus, a possible explanation for the good performance of *ℓ*_2_-penalization is that it does not need an optimal choice of regions, perfectly aligned with the underlying functional modules.

### 3.5. Effect of the dataset size and heterogeneity

While not a property of the data-processing pipeline itself, a large number of training subjects is the most important factor of success for prediction. Indeed, we observe that prediction results improve with the number of subjects, even for large number of subjects (Figure 3).

Despite that increase, we note that prediction on the largest subsample (subsample #1) gives lower scores than prediction on subsample #2, which may be due to the variance introduced by small sites. We also note that the importance of the choice between each option of the pipeline decreases with the number of subjects (Figure A8).

### 3.6. Effect of the number of regions

In order to compare models of similar complexity, the previous experiments were restricted to 84 regions as in [49]. However, this choice is based on previous studies and may not be the optimal choice for our problem. To validate this aspect of the study, we also varied the number of regions and analyzed its impact on the classification results in Figure 5, similarly to Figure 3. In order to avoid computational cost^12^, we explored only two atlases: the data-driven Ward’s hierarchical clustering and spectral clustering atlases computed in [43].

**Figure 5:**
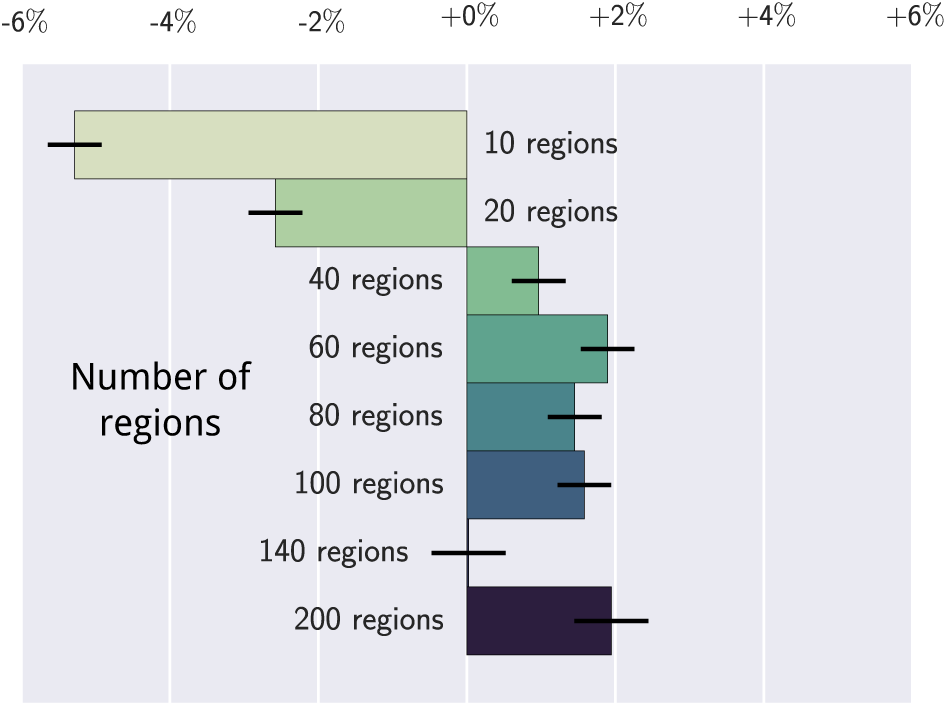
Impact of the number of regions on prediction. each bar indicates the impact of the number of regions on the prediction accuracy relative to the mean of the prediction. These values are coefficients in a linear model explaining the best classification scores as function of the number of regions. Error bars give the 95% confidence interval, computed by a full factorial ANOVA. Atlases containing more than 40 ROIs give better results in all settings. Results per subsamples are presented in the appendices –Figure A9.

Across all settings, we distinguish 3 different regimes. First, below 20 ROIs, performance is bad. Indeed very large regions average out signals of different functional areas. Above 140 ROIs, the results seem to be unstable, though this trend is less clear in intra-site prediction. Finally, the best results are between 40 and 100 ROIs. Our choice of 84 ROIs is within the range of best performing models.

### 3.7. Inspecting discriminative connections

A connection between two regions is considered discriminative if its value helps diagnosing ASD from TC. To understand what is driving the best prediction pipeline, we extract its most discriminative connections as described in section 2.7. Given that the 10-fold cross-validation yields 10 different atlases, we start by building a consensus atlas by taking all the regions with a DICE similarity score above 0.9. We obtain an atlas composed of 37 regions (see Figure 6). These regions and the discriminative connections form the neurophenotypes. Note that discriminative connections should be interpreted with caution. Indeed, our study covers a wide range of age and diagnostic criteria. Additionally, predictive models cannot be interpreted as edge-level tests.

**Figure 6:**
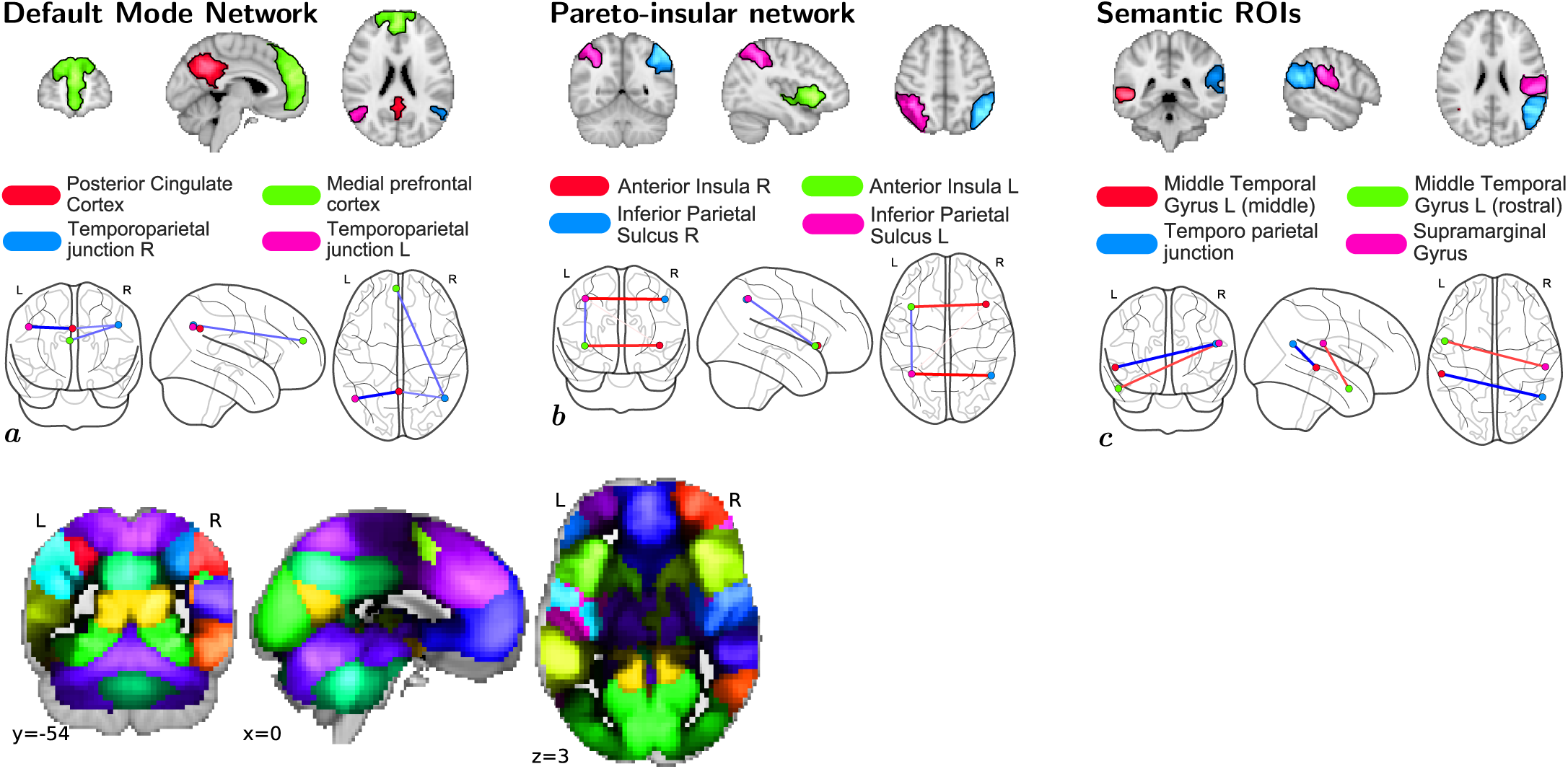
Significant non-zero connections in the predictive biomarkers distinguishing controls from ASD patients. Red connections are stronger in controls and blue connections are stronger in ASD patients. Subfigures *a*, *b*, and *c* reported intranetwork difference. On the left is the consensus atlas obtained by selecting regions consistently extracted on 10 subsets of ABIDE. Colors are randomly assigned. Results obtained on other networks are shown in the appendices –Figure A7.

*Default Mode Network (Figure 6.a).* We observe a lower connectivity between left and right temporo-parietal junctions. Both hypoconnectivity between regions on adolescent and adult patients [59, 60] and hyperconnectivity in children [61] have been reported. Our findings are concordant with [62] that observed lower fronto-parietal connectivity and stronger connectivity between temporal ROIs.

*Pareto-insular network (Figure 6.b).* We observe inter-hemispheric hypoconnectivity between left and right anterior insulae, and left and right inferior parietal lobes. Those regions are part of a frontoparietal control network related to learning and memory [63]. Such hypo-connectivity has already been widely evoked in ASD [64, 65] and previously found in the ABIDE dataset [13, 66]. These ROIs also match anatomical regions of increased cortical density in ASD [67].

*Semantic ROIs (Figure 6.c).* We observe a more complicated pattern in ROIs related to language. Connectivity is stronger between the right supramarginal gyrus and the left Middle Temporal Gyrus (MTG) while a lower connectivity is observed between the right MTG and the left temporo-parietal junction. Differences of laterality in language networks implicating the middle temporal gyri have already been observed [68]. Although this symptom is often considered as decorrelated from ASD, we find that these regions are relevant for diagnosis.

## 4. Discussion

We studied pipelines that extract neurophenotypes from aggregate R-fMRI datasets through the following steps: 1) region definition from R-fMRI, 2) extraction of regions activity time series 3) estimation of functional interactions between regions, and 4) construction of a discriminant model for brain-based diagnostic classification. The proposed pipelines can be built with the Nilearn neuroimaging-analysis software and the atlas computed with MSDL is available for download ^13^. We have demonstrated that they can successfully predict diagnostic status across new acquisition sites in a real-world situation, a large Autism database. This validation with a leave-onesite-out cross-validation strategy reduces the risk of overfitting data from a single site, as opposed to the commonly used leave-one-subject-out validation approach [53]. It enabled us to measure the impact of methodological choices on prediction. In the ABIDE dataset, this approach classified the label of autism versus control with an accuracy of 67% – a rate superior to prior work using the larger ABIDE sample.

The heterogeneity of weakly-controlled datasets formed by aggregation of multiple sites poses great challenges to develop brain-based classifiers for psychiatric illnesses. Among those most commonly cited are: *i)* the many sources of uncontrolled variation that can arise across studies and sites (*e. g.* scanner type, pulse sequence, recruitment strategies, sample composition), *ii)* the potential for developing classifiers that are reflective of only the largest sites included, and *iii)* the danger of extracting biomarkers that will be useful only on sites included in the training sample. Counter to these concerns, our results demonstrated the feasibility of using weakly-controlled heterogeneous datasets to identify imaging biomarkers that are robust to differences among sites. Indeed, our prediction accuracy of 67% is the highest reported to-date on the whole ABIDE dataset (compared to 60% in [53]), though better prediction has been reported on curated more homogeneous subsets [13, 54]. Importantly, we found that increasing sample size helped tackling heterogeneity as intersite prediction accuracy approached that of intra-site prediction. This undoubtedly increases the perceived value that can be derived from aggregate datasets, such as the ABIDE and the ADHD-200 [16].

Our results suggest directions to improve prediction accuracy. First, more data is needed to optimize the classifier. Indeed, our experiments show that with a training set of 690 participants aggregated across multiple sites (80% of 871 available subjects), the pipelines have not reached their optimal performance. Second, a broader diversity of data will be needed to more definitively assess prediction for sites not included in the training set. Both of these needs motivate more data sharing.

Methodologically, our systematic comparison of choices in the pipeline steps can give indications to establish standard processing pipelines. Given that processing pipelines differ greatly between studies [28], comparing data-analysis methods is challenging. On the ABIDE dataset, we found that a good choice of processing steps can strongly improve prediction accuracy. In particular, the choice of regions is most important. We found that extracting these regions with MSDL gives best results. However, the benefit of this method is not always clear cut – on small datasets, references atlases are also good performers. For the rest of the pipeline, tangent embedding and *ℓ*_2_-regularized classifier are overall the best choice to maximize prediction accuracy.

While the primary goals of this study were methodological, it is worth noting that biomarkers identified were largely consistent with the current understanding of the neural basis of ASD. The most informative features to predict ASD concentrated in the intrinsic functional connectivity of three main functional systems: the defaultmode, parieto-insular, and language networks, previously involved in ASD [69, 70, 71]. Specifically, decreased homotopic connectivity between temporo-parietal junction, insula and inferior parietal lobes appeared to characterize ASD. Decreases in homotopic functional connectivity have been previously reported in R-fMRI studies with moderately [64, 72] or large samples such as ABIDE [25]. Here our findings point toward a set of inter-hemispheric connections involving heteromodal association cortex subserving social cognition (*i. e.* temporo-parietal junction [73]), and cognitive control (anterior insula and right inferior parietal cortex [74]), commonly affected in ASD [64, 65, 13, 66]. Limitations of our biomarkers may arise from the heterogeneity of ASD as a neuropsychiatric disorder [75].

Beyond forming a neural correlate of the disorder, predictive biomarkers can help define diagnosis subgroups – the next logical endeavor in classification studies [76].

As a psychiatric study, the present work has a number of limitations. First, although composed of samples from geographically distinct sites, the representativeness of the ABIDE sample has not been established. As such, the features identified may be biased and necessitate further replication; though their consistency with prior findings alleviates this concern to some degree. Further work calls for characterizing this prediction pipeline on more participants (*e. g.* ABIDE II) and different pathologies (*e.g.* ADHD-200 dataset). Second, while the present work primarily relied on prediction accuracy to assess the classifiers derived, a variety of alternative measures exist (*e.g.* sensitivity, specificity, J-statistic). This decision was largely motivated to facilitate comparison of our findings with those from prior studies using ABIDE, which most frequently used prediction accuracy. Depending on the specific application for a classifier, the prediction metric to optimize may vary. For example screening tools would need to favor sensitivity over accuracy or specificity; in contrast with medical diagnostic tests that require specificity over sensitivity or accuracy [4]. The versatility of the pipeline studied allows us to maximize either of these scores depending on the experimenter’s needs. Finally, while the present work focused on the ABIDE dataset, it is likely that its findings regarding optimal pipeline decisions will carry over to other aggregate samples, as well as more homogeneous samples. With that said, future applications will be required to verify this point.

In sum, we experimented with a large number of different pipelines and parametrizations on the task of predicting ASD diagnosis on the ABIDE dataset. From an extensive study of the results, we have found that R-fMRI from a large amount of participants sampled from many different sites could lead to functional-connectivity biomarkers of ASD that are robust, including to inter-site variability. Their predictive power markedly increases with the number of participants included. This increase holds promise for optimal biomarkers forming good probes of disorders.

## Acknowledgments.

We acknowledge funding from the NiConnect project and the SUBSample project from the DIGITEO Institute, France. The effort of Adriana Di Martino was partially supported by the NIMH grant 1R21MH107045-01A1. Finally, we thank the anonymous reviewers for their feedback, as well as all sites and investigators who have worked to share their data through ABIDE.

## Footnotes

1 See http://fcon_1000.projects.nitrc.org/indi/abide/ for specific information

2 http://preprocessed-connectomes-project.github.io/abide

3 For Harvard-Oxford and Yeo reference atlases, selecting 84 ROIs yields slightly better results than using the full version as shown in Figure A2. This is probably due to the fact that these additional regions are smaller and less important regions, hence they induce spurious correlations in the connectivity matrices.

4 We identify these ROIs using an approach similar to FSL FIX (FMRIB’s ICA-based Xnoiseifier). FIX extracts spatial and temporal descriptors from matrix decomposition results and uses a classifier trained on manually labeled components to determine if they correspond to noise.

5 For the sake of simplicity, we report only the 3 main methods here. However, we also investigated random forests and Gaussian naive Bayes, that gave low prediction performance. Feature selection also led to worse results, both with univariate ANOVA screening and with sparse models such as Lasso or SVC-*ℓ*_1_ see Figure A4 for more details. Logistic regression gave results similar to SVC.

6 Software versions: python 2.7, scikit-learn 0.17.1, scipy 0.14.0, numpy 1.9.0, nilearn 0.1.5.

7 The list of subjects used for each cross-validation fold are available at https://team.inria.fr/parietal/files/2016/04/cv_abide.zip

8 Note that the criterion used to select the data-point used for the post-hoc analysis, i. e. the 10% best performing, is independent from the factors studied in this analysis, as we select the 10% best performing for each value of these factors. Hence, the danger of inating effects by a selection [56] does not apply to the post hoc analysis.

9 Despite the optimizations, the whole study presented here is very computationally costly. Indeed, the nested cross-validation, the various pipeline options, and the various subsets, would lead to a computation time of 10 years on a computer with 8 cores and 32GB RAM.We used a computer grid to reduce it to two months.

10 Chance is calculated by taking the highest score obtained by a dummy classifier that predicts random labels following the distribution of the two classes.

11 Note that we experimented with a variety of more sophisticated covariance estimators, including GraphLasso, though without improved results.

12 As MSDL is much more computationally expensive than Ward, we have not explored the effect of modifying of the number of regions extracted with MSDL. However, we expect that the optimal number of regions is not finely dependent on the region-extraction method.

13 http://team.inria.fr/parietal/files/2015/07/MSDL_ABIDE.zip

